# Ketone-body receptor GPR109A suppresses hepatic inflammation via gut–liver axis regulation

**DOI:** 10.1101/2025.08.21.671439

**Authors:** Akari Nishida, Shota Nishikawa, River Budau, Mayu Yamano, Ryuji Ohue-Kitano, Takako Ikeda, Nobuo Sasaki, Ikuo Kimura

## Abstract

The ketogenic diet (KD) promotes ketone body synthesis and has been used as an effective treatment for disorders such as epilepsy. Although elevated ketone bodies, including β-hydroxybutyrate (βHB) and acetoacetate, are thought to meditate the beneficial effects of the KD, the mechanisms underlying their metabolic actions remain incompletely understood. In this study, we focused on GPR109A, a receptor for βHB with an unclear role in metabolic homeostasis. We employed KD and fasting models to examine metabolic changes under two distinct ketogenic conditions. Under KD conditions, *Gpr109a^-/-^* mice exhibited increased hepatic lipid accumulation, and subsequent hepatic inflammation and fibrosis. However, *Gpr109a* deletion did not exacerbate hepatic lipid accumulation or inflammation during short-term fasting, suggesting that GPR109A-mediated liver protection is specific to KD-induced metabolic stress rather than under fasting conditions. Mechanistic analysis revealed that GPR109A protects the liver from inflammation by maintaining intestinal barrier integrity. These findings highlight the novel protective mechanism of GPR109A, via the gut–liver axis, to sustain metabolic homeostasis during the KD. This study provides valuable insights into the physiological effects of ketone bodies.

## 1 INTRODUCTION

Mammals ingest and efficiently convert exogenous energy sources into metabolic products. Among these, ketone bodies are produced during conditions such as fasting and diabetes, in which glucose utilization is limited. Ketogenesis increases in proportion to total fat oxidation. After the β-oxidation of fatty acids, acetyl-CoA is converted into acetoacetyl-CoA and further converted into HMG-CoA by HMGCS2, the key enzyme for ketone synthesis. HMG-CoA is converted into acetoacetate, a ketone body, by HMGCL and subsequently converted into β-hydroxybutyrate (βHB), another ketone body, through a continuous enzymatic process involving BDH1. Once taken up by extrahepatic tissues, including the brain and heart, βHB is oxidized by BDH1 and SCOT and enters the tricarboxylic acid cycle, ultimately leading to the production of ATP (1).

Although ketone body production is typically restricted to periods of glucose depletion, certain dietary interventions actively promote ketone body synthesis. Fasting was first recorded as a therapeutic measure for treating seizures in the Hippocratic corpus (in the 5^th^ century BCE), and from the early 20^th^ century on, clinical trials have demonstrated that the ketogenic environment contributes to improvements in epilepsy symptoms. Since then, ketogenic diets (KDs) have been reported to be beneficial for various disorders, including epilepsy, obesity, cancer, and respiratory diseases (2, 3). The classic KD, in which 90% of the caloric intake is derived from lipids, is widely used in clinical practice. Furthermore, since 1971, medium-chain triglycerides (MCTs) have been found to enhance ketone body production, and modified Atkins diets and low-glycemic index therapies have been developed as alternatives (4–6). Although the KD is effective for certain disorders, it can also increase LDL-C and total cholesterol levels and exacerbate fatty liver due to its high lipid content (7–10). Nevertheless, how the body senses ketone bodies and what physiological responses are elicited under ketogenic conditions remain poorly understood at the mechanistic level.

Regarding their metabolic effects, ketone bodies serve not only as an energy source but also as signaling molecules through G-protein–coupled receptors (GPCRs). We have previously reported that βHB serves as an antagonist for GPR41/FFAR3, and βHB-mediated signaling through GPR41 suppresses sympathetic nervous system activity, thereby modulating energy expenditure (11). In addition, we identified GPR43/FFAR2 as a receptor for acetoacetate and demonstrated that the acetoacetate–GPR43 axis promotes lipolysis (12). These findings support an important role of ketone bodies as signaling molecules in the regulation of energy metabolism.

GPR109A/HCAR2 was initially identified as a receptor for nicotinic acid and butyrate and was later recognized as a receptor for βHB in 2005 (13). GPR109A, coupled with Gα/io, is expressed in immune cells, adipose tissue, and colonic epithelial cells (14). Although activation of GPR109A on Langerhans cells, immune cells in the skin, mediates flushing—the side effect of nicotinic acid—via β-arrestin signaling (15), GPR109A also plays beneficial roles in immune functions. Previous studies have shown that GPR109A mediates a microglial protective response, which consequently attenuates amyloid-induced pathology (16). Furthermore, it has been reported that butyrate suppresses colonic inflammation by inducing the differentiation of Treg cells and IL-10–producing T cells via GPR109A (17). In addition, intraperitoneal administration of nicotinic acid has been shown to inhibit lipolysis (18), suggesting that GPR109A plays a critical role in metabolic regulation. Collectively, these findings have drawn attention to GPR109A as a potential therapeutic target for immune and metabolic disorders. However, considering that physiological concentrations of nicotinic acid do not activate GPR109A and that butyrate-mediated activation occurs only under specific conditions within the intestinal tract(19, 20), the physiological role of GPR109A is not yet fully understood. Additionally, in vivo studies investigating βHB–GPR109A signaling remain limited. In this study, we investigated the physiological role of GPR109A under ketogenic conditions using *Gpr109a*-deficient mice.

## 2 RESULTS

### 2.1 Ketogenic diet induces hepatic inflammation in *Gpr109a*-deficient mice

To investigate the role of GPR109A in a ketogenic environment, wild-type (WT) and *Gpr109a^-/-^* mice were subjected to a KD predominantly composed of fat and a small amount of carbohydrates (**Supplementary Figure 1, Supplementary Table 1**). Plasma βHB concentrations were significantly elevated in both KD-fed mice (**Figure 1A**). As previously reported (12), body weight gain was suppressed in mice fed the KD compared with those fed the normal diet (ND) **(Supplementary Figure 2A)**. Under these conditions, no significant differences in body weight were observed between wild-type and *Gpr109a⁻/⁻* mice **(Figure 1B)**. Adipose tissue weight was also comparable between the two groups **(Figure 1C)**. In contrast, liver weight was significantly higher in *Gpr109a⁻/⁻* mice than in WT mice **(Figure 1D)**. To evaluate hepatic lipid accumulation, we measured hepatic triglyceride levels. KD feeding, which represents an ultra-high-fat dietary intervention, led to an increase in hepatic triglyceride content in both WT and *Gpr109a⁻/⁻*mice. Notably, hepatic triglyceride levels were significantly higher in *Gpr109a⁻/⁻* mice compared with WT mice (**Figure 1E**). Consistent with these biochemical findings, histological examination of the liver under KD conditions demonstrated a more pronounced accumulation of lipid droplets in *Gpr109a⁻/⁻* mice than in their WT counterparts (**Figure 1F**). However, hepatic total cholesterol levels were similar between WT and *Gpr109a^-/-^* mice (**Figure 1E**). Similar to adipose tissue weight (**Figure 1C**), there were no significant differences in blood glucose levels and circulating lipids between the two groups (**Supplementary Figure 2B-E**). Furthermore, hepatic mRNA profiling in KD-fed mice revealed that several *de novo* lipogenesis–related genes, particularly those associated with fatty acid synthesis, were upregulated in *Gpr109a^⁻/⁻^* mice, whereas genes related to lipid uptake showed no significant differences compared with WT mice (**Figure 1G**). These results suggest that GPR109A deficiency promotes hepatic lipid accumulation through increased hepatic fatty acid synthesis, independent of adipose tissue lipolysis.

**FIGURE 1.**
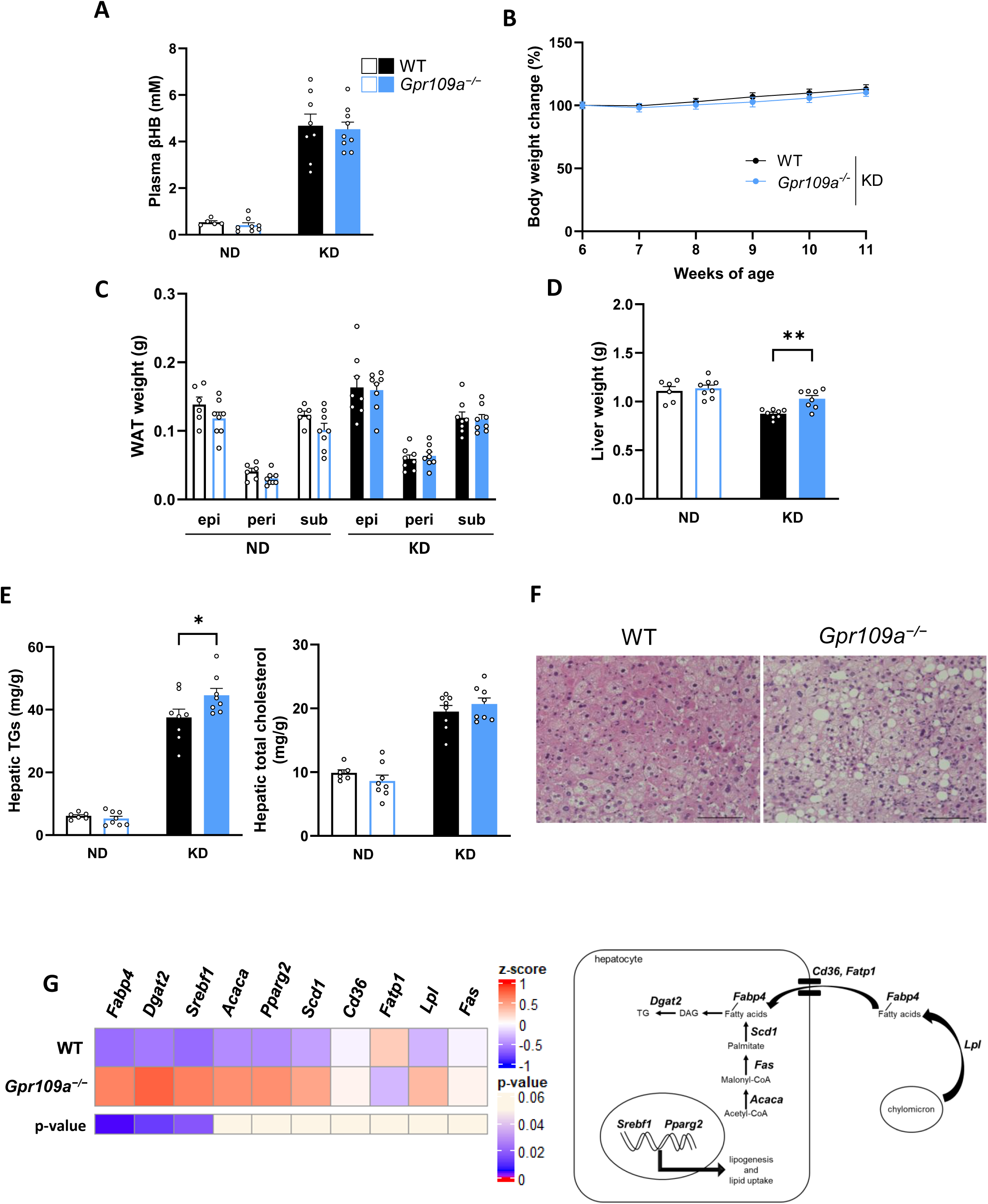
*Gpr109a^−/−^* mice show increased hepatic lipid accumulation under KD feeding. **(A)** Plasma β-Hydroxybutyrate of mice fed a normal diet (ND) or ketogenic diet (KD) for 6 weeks (n = 5–9). **(B)** Body weight changes in wild-type (WT) and *Gpr109a^−/−^* mice fed the KD (n = 8). **(C)** Adipose tissue weight (n = 6–8). **(D)** Liver weight (n = 6–8). **(E)** Hepatic triglycerides and total cholesterol (n = 6–8). **(F)** HE-stained liver samples from WT and *Gpr109a^−/−^*mice fed the KD. Scale bar = 100 µm. epi, epididymal adipose tissues; peri, perirenal adipose tissues; sub, subcutaneous adipose tissues. **(G)** Heatmap showing the expression of *de novo* lipogenesis- and lipolysis-related genes in the liver of WT and *Gpr109a⁻/⁻* mice fed the ND or KD (n = 6–8). ** P < 0.01; * P < 0.05, two-way ANOVA with Sidak’s multiple comparisons test: A, B, C, D, E, Mann–Whitney U test: G. All data are presented as the mean ± SEM.

### 2.2 GPR109A modulates immune cell recruitment to protect against hepatic inflammation

We measured representative inflammatory (*Tnf*) and fibrotic (*Col1a*) markers to evaluate hepatic inflammation, which is associated with hepatic lipid accumulation. While KD feeding increased the levels of these markers in both WT and *Gpr109a^-/-^* mice, their expression was significantly higher in *Gpr109a^-/-^* mice **(Figure 2A)**. Immunostaining of the liver in KD-fed mice also revealed increased levels of the macrophage marker F4/80 and the fibrosis marker α-smooth muscle actin (α-SMA) in the liver of *Gpr109a^-/-^* mice **(Figure 2B)**. We investigated the molecular mechanisms by which GPR109A protects the liver from inflammation under the KD condition. GPR109A is expressed in immune cells (17), and our flow cytometry analysis confirmed its high expression in bone marrow-derived monocytes and macrophages, but not in tissue-resident Kupffer cells (**Figure 2C**). Next, we evaluated the potential anti-inflammatory effects of βHB using RAW264.7 macrophage-like cells. Treatment with capric acid (C10), previously reported to have anti-inflammatory effects, significantly reduced *Tnf* expression (21). In contrast, βHB did not suppress *Tnf* expression **(Figure 2D)**. Similarly, MK6892, a potent synthetic agonist of GPR109A, did not reduce *Tnf* expression **(Figure 2D)**. These results indicate that GPR109A is unlikely to directly regulate *Tnf* expression in macrophages. In *Gpr109a^⁻/⁻^* mice under KD conditions, the expression of *Adgre1*, a macrophage marker, was elevated **(Figure 2E)**. Furthermore, the expression of *Mcp1* (CCL2), a chemokine that promotes monocyte chemotaxis, was significantly increased in *Gpr109a⁻/⁻* mice **(Figure 2F)**. Taken together, these findings suggest that, under diet-induced ketogenic conditions, the liver of *Gpr109a^⁻/⁻^* mice exhibits a microenvironment that is more permissive to macrophage recruitment.

**FIGURE 2.**
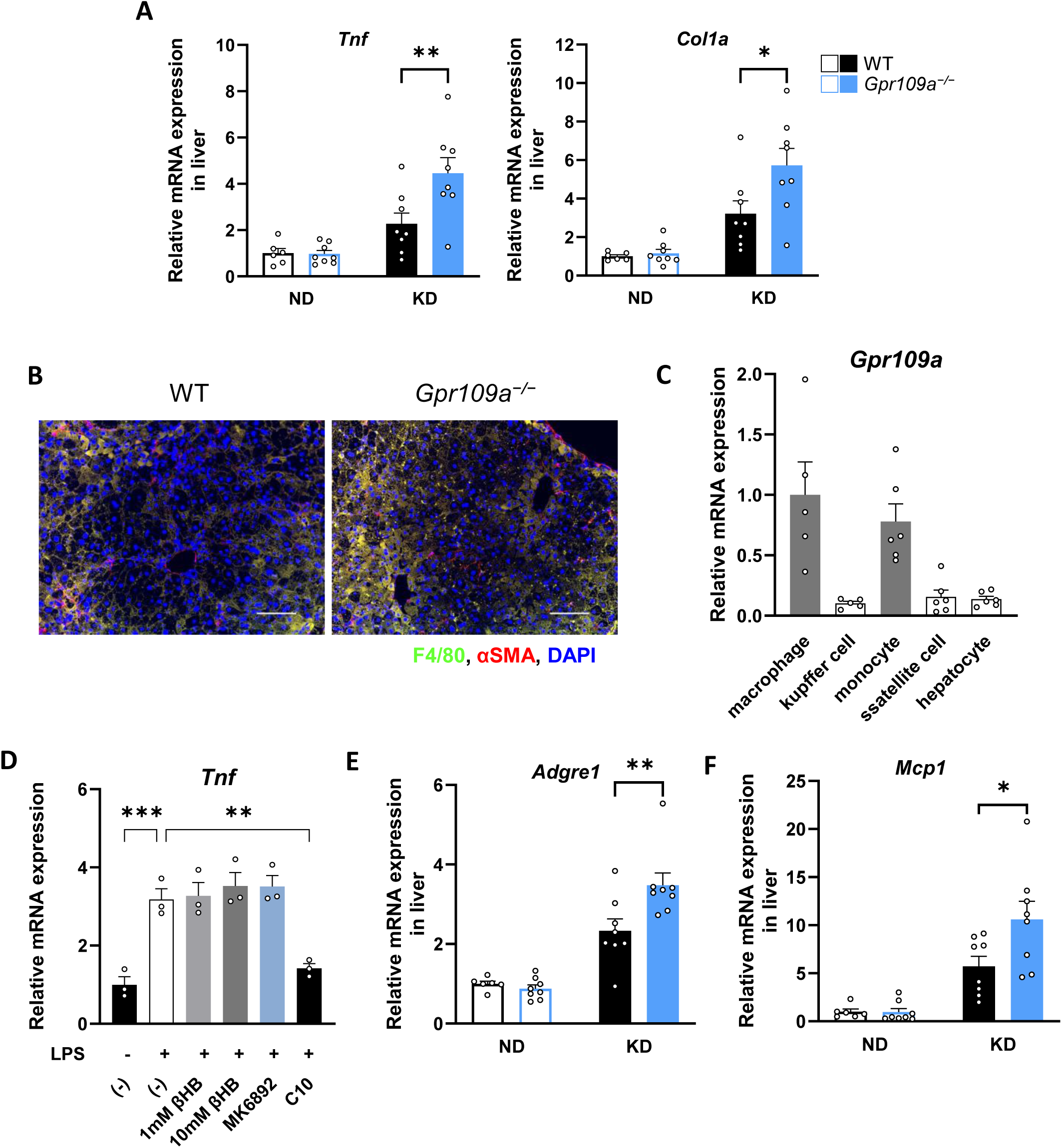
GPR109A suppresses hepatic inflammation. **(A)** mRNA expression levels of inflammation- and steatosis-related genes in the liver in WT and *Gpr109a^−/−^* mice fed the KD (n = 6–8). **(B)** Inflammation levels in the liver. Scale bar = 100 µm. **(C)** *Gpr109a* expression in hepatic macrophages, bone marrow (BM)-derived monocytes, Kupffer cells, hepatic stellate cells, and hepatocytes isolated from WT mice fed the ND (n = 5–6). **(D)** Anti-inflammatory effect of β-Hydroxybutyrate—or MK6892—stimulated GPR109A in RAW264.7 cells (n = 3). **(E)** mRNA expression levels of a macrophage marker in the liver (n = 6–8). **(F)** mRNA expression levels of the monocyte chemoattractant protein in the liver (n = 6–8). *** P < 0.001; ** P < 0.01; * P < 0.05, two-way ANOVA with Sidak’s multiple comparisons test: A, E, F; Dunnett’s test: D. All data are presented as the mean ± SEM.

### 2.3 GPR109A does not contribute to the hepatoprotective effect during short-term fasting

In our model, KD-driven ketosis (high-fat intake) induced hepatic lipid accumulation and subsequent inflammation. Under this condition, GPR109A plays a role in liver protection. Therefore, we next performed a fasting experiment as an acute ketogenic condition associated with energy shortage. Consistent with the KD condition, plasma βHB levels increased after 24 h fasting, regardless of genotype (**Figure 3A)**. In the fasting condition, *Gpr109a^-/-^* mice exhibited body weight reductions comparable to those of WT mice (**Figure 3B**). Unexpectedly, no significant difference in liver weight was observed between the two groups under fasting conditions (**Fig. 3C**). Moreover, although hepatic triglyceride and total cholesterol levels increased in both genotypes following fasting, no significant difference was detected between the two groups (**Figure 3D**). To further investigate these findings, we performed hepatic gene expression analysis. Unlike under KD conditions, fasting itself did not induce hepatic inflammation, fibrosis, or macrophage infiltration. Accordingly, no statistical difference was found in the expression of hepatic inflammation, fibrosis, and macrophage markers between WT and *Gpr109a^⁻/⁻^* mice (**Figure 3E**). No significant difference was observed in adipose tissue weight, blood glucose levels, or circulating lipids (**Supplementary Figure 3A-E**). These results suggest that GPR109A does not contribute to liver protection during short-term fasting. Instead, its protective effects may be more prominent under specific ketogenic conditions, where it appears to counteract the detrimental effects of a lipid-rich diet.

**FIGURE 3.**
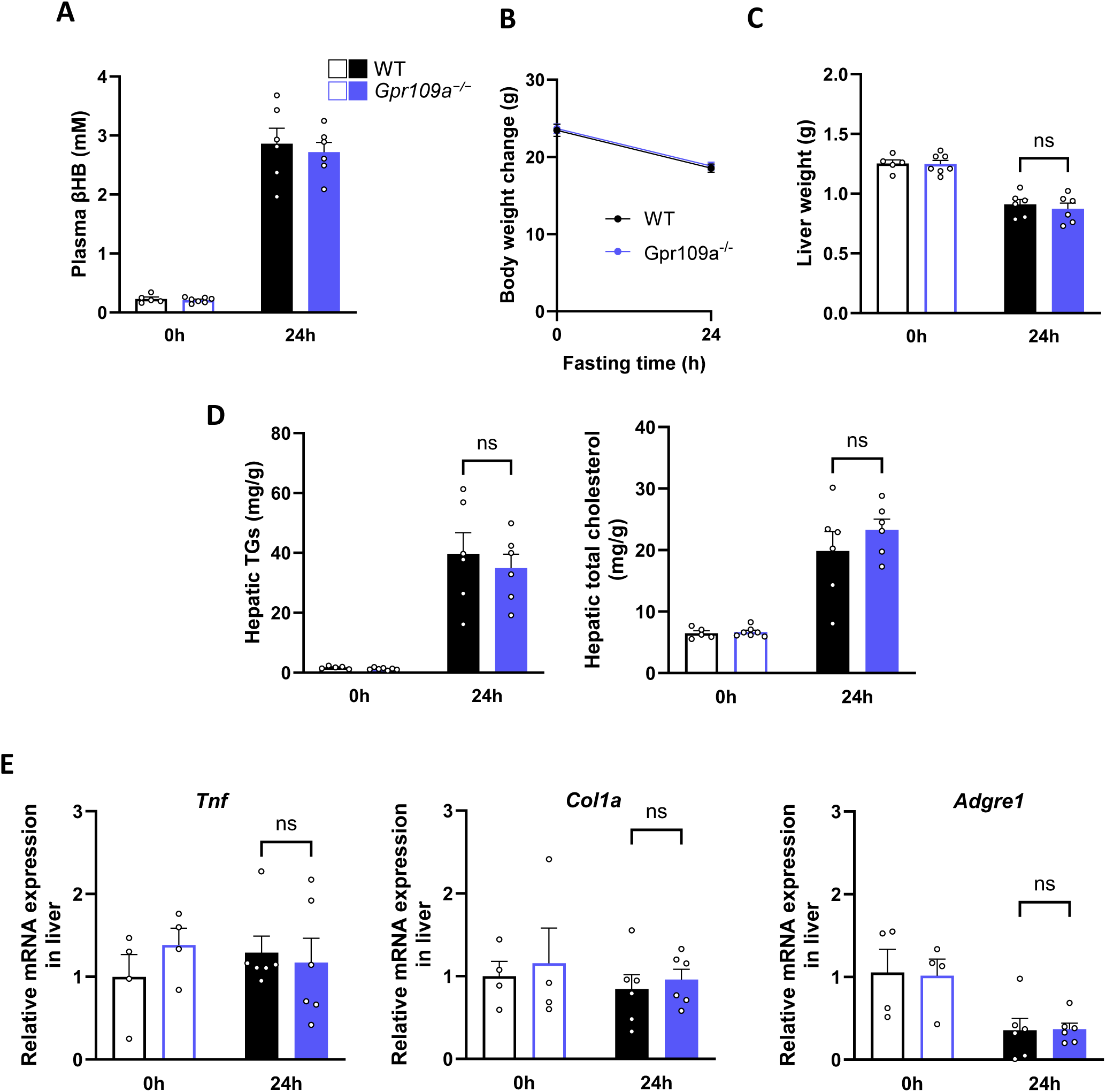
*Gpr109a* deficiency does not cause metabolic dysfunction under fasting-induced ketogenic conditions. **(A)** Plasma β-Hydroxybutyrate after 0 and 24 h of fasting (n = 5–7). **(B)** Body weight changes in WT and *Gpr109a^−/−^* mice after 24 h of fasting (n = 6). **(C)** Liver weight (n = 5–7). **(D)** Hepatic triglycerides and total cholesterol (n = 5–7). **(E)** mRNA expression levels of inflammation- and steatosis-related genes and a macrophage marker in the liver in WT and *Gpr109a^−/−^* mice after 0 or 24 h of fasting (n = 4–6). two-way ANOVA with Sidak’s multiple comparisons test: A–E. All data are presented as the mean ± SEM.

### 2.4 GPR109A modulates intestinal tight junctions and contributes to liver protection under diet-induced ketogenic conditions

Based on the aforementioned results, we demonstrated that the hepatic lipid accumulation and inflammation observed in *Gpr109a^-/-^*mice were induced only under specific ketogenic conditions. To investigate this further, we performed a detailed analysis of mRNA expression in the livers of mice fed the KD and found that some TLR4 signaling–related genes were upregulated in *Gpr109a^-/-^* mice (**Figure 4A**), indicating that the influx of PAMPs (Pathogen-Associated Molecular Patterns) could cause hepatic inflammation. GPR109A is known to regulate intestinal tight junctions (TJs) in response to butyrate-stimulation (22, 23). Therefore, we measured the expression of TJ markers in the colon and found reduced expression of several TJ markers, particularly *Cldn3*, in *Gpr109a^-/-^*mice compared with WT mice under diet-induced ketogenic conditions (**Figure 4B**). Immunostaining of the colon further confirmed decreased Claudin3 levels in *Gpr109a^⁻/⁻^* mice fed the KD diet **(Figure 4C)**. These results suggest that high dietary lipid load can disrupt intestinal barrier integrity (24), whereas βHB produced during KD feeding mitigate this barrier dysfunction via GPR109A. In contrast, no such reduction in TJ marker expression was observed under fasting conditions (**Figure 4D**). Interestingly, the decrease in TJ marker expression was not observed in the small intestine of *Gpr109a^-/-^* mice fed the KD, where the expression of GPR109A was much lower than that in the colon (**Supplementary Figure 4A**). Moreover, depletion of gut microbiota by antibiotic treatment abolished the enhanced hepatic inflammation observed in *Gpr109a^⁻/⁻^* mice under KD feeding (**Figure 4E, Supplementary Figure 4B**). These results indicate that under diet-induced ketogenic conditions, βHB–GPR109A signaling may contribute to the maintenance of TJ in the colon, thereby protecting hepatic function.

**FIGURE 4.**
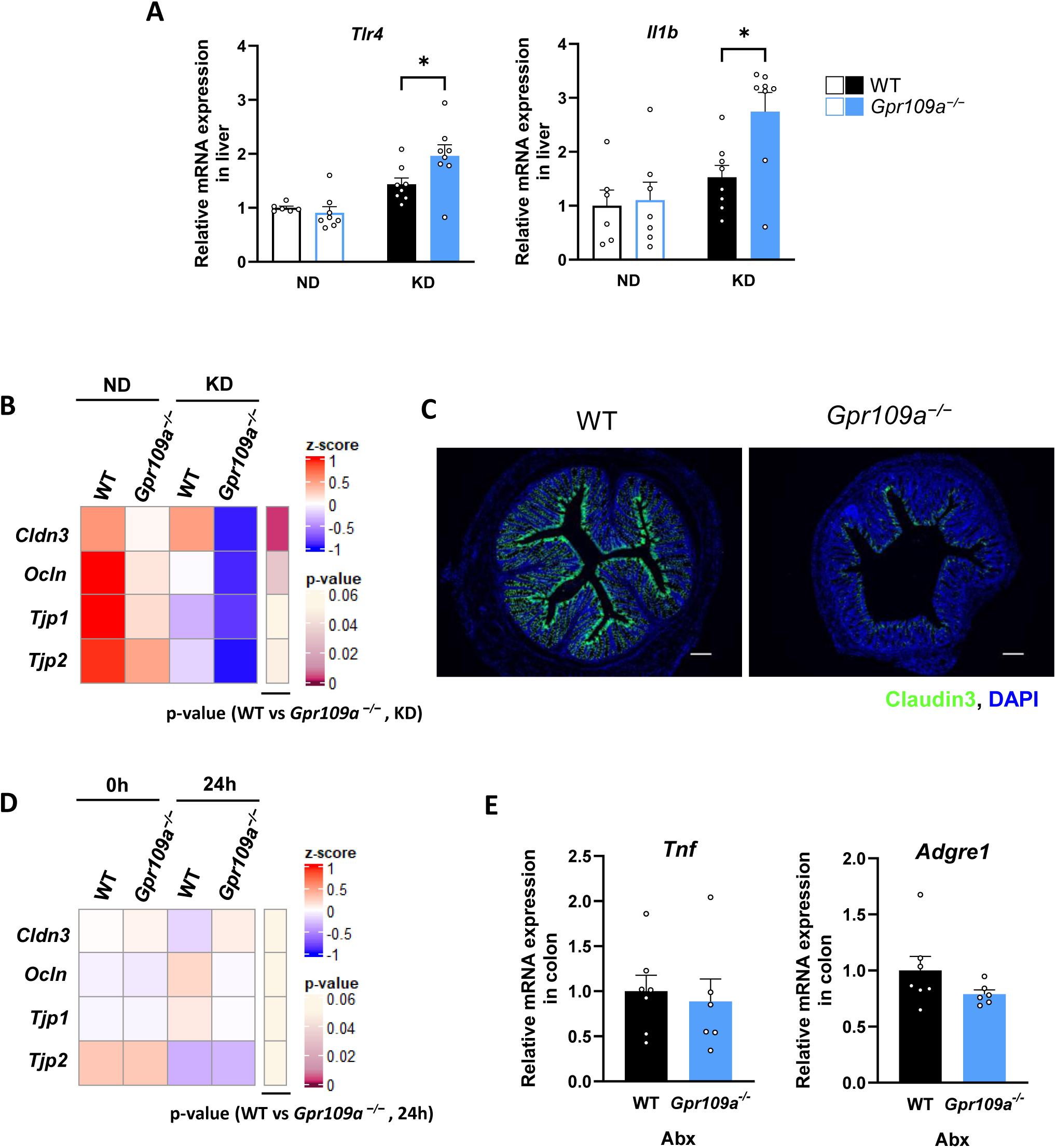
GPR109A protects tight junctions in the colon under diet-induced ketogenic conditions. **(A)** mRNA expression levels of lipopolysaccharide (LPS)-related genes in the liver in WT and *Gpr109a^−/−^* mice fed the KD (n = 6–8). (**B**) Heatmap showing the expression of tight junction-related genes in the colon of WT and *Gpr109a^−/−^* mice fed the ND or KD (n = 6–8). **(C)** Tight junction protein levels in the colon under KD conditions. Scale bar = 100 µm. **(D)** Heatmap showing the expression of tight junction-related genes in the colon of WT and *Gpr109a^−/−^* mice under 0 or 24 h fasting-induced ketosis (n = 6–7). **(E)** mRNA expression levels of inflammation- and steatosis-related genes in the liver of WT and *Gpr109a^−/−^* mice fed the KD and treated with an antibiotic cocktail (n = 6–7). ** P < 0.01, * P < 0.05, two-way ANOVA with Sidak’s multiple comparisons test: A, B, D; Student’s *t*-test: E. All data are presented as the mean ± SEM.

### 2.5 Colonic ketogenesis and cellular distribution of GPR109A suggest a paracrine β-hydroxybutyrate signaling axis in the colon

Previous studies have demonstrated that HMGCS2, the rate-limiting enzyme in ketogenesis, is highly expressed not only in the liver but also in the colon (25, 26). To further investigate colonic ketogenesis and the role of GPR109A in TJ regulation, we examined *Hmgcs2* expression in organs known to express this enzyme, including the colon, and measured colonic βHB levels under diet-induced ketogenic conditions. The results revealed that high *Hmgcs2* expression in the colon led to a marked increase in colonic βHB levels in mice fed the KD (**Figure 5A, B**). Next, to identify the cellular sources of *Gpr109a* in the colon, we reanalyzed a public single-cell RNA-seq dataset (27). Consistent with previous reports, *Gpr109a* expression was detected in epithelial cells and macrophages, but its expression was more pronounced in fibroblasts **(Figure 6A, B, Supplementary Figure 5)**. Moreover, *Gpr109a* expression was confirmed in both isolated fibroblast cell cultures and epithelial organoids **(Figure 6C)**. Taken together, these results suggest that colonic βHB may act within the intestinal microenvironment to activate GPR109A in a paracrine manner, thereby contributing to the intestinal homeostasis.

**FIGURE 5.**
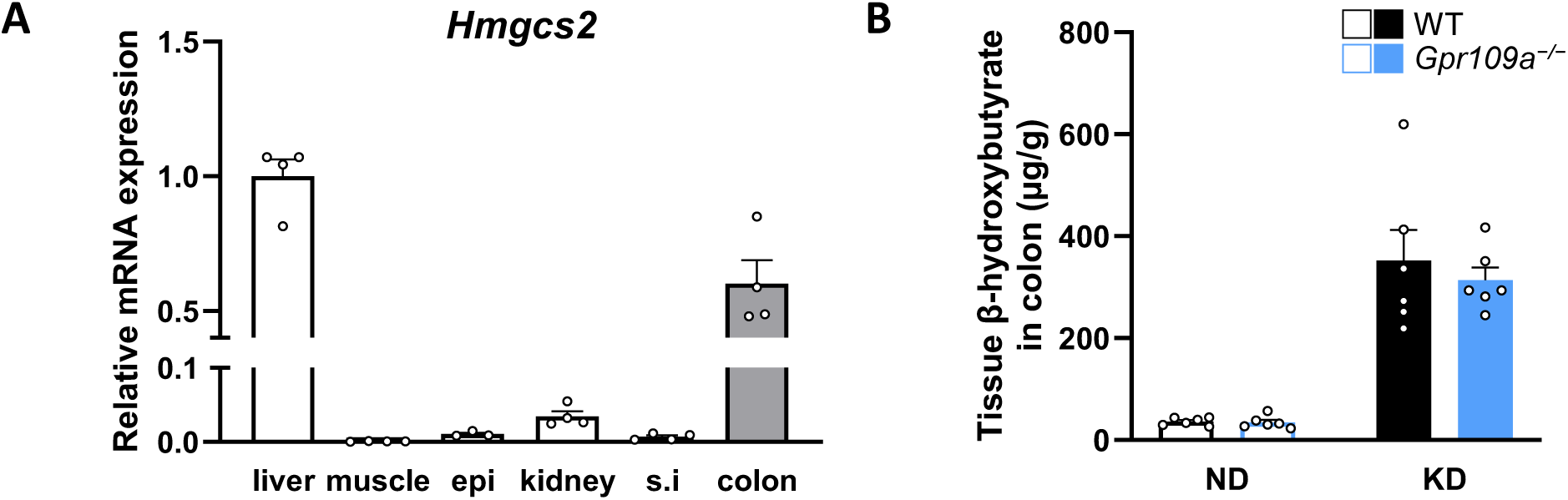
The colon contributes to ketone body production. **(A)** Expression of *Hmgcs2* in the liver, muscle, epididymal adipose tissue, kidney, small intestine, and colon of WT mice (n = 3–4). **(B)** β-Hydroxybutyrate concentration in the colon tissue of mice fed the ND or KD for 6 weeks (n = 6).

**FIGURE 6.**
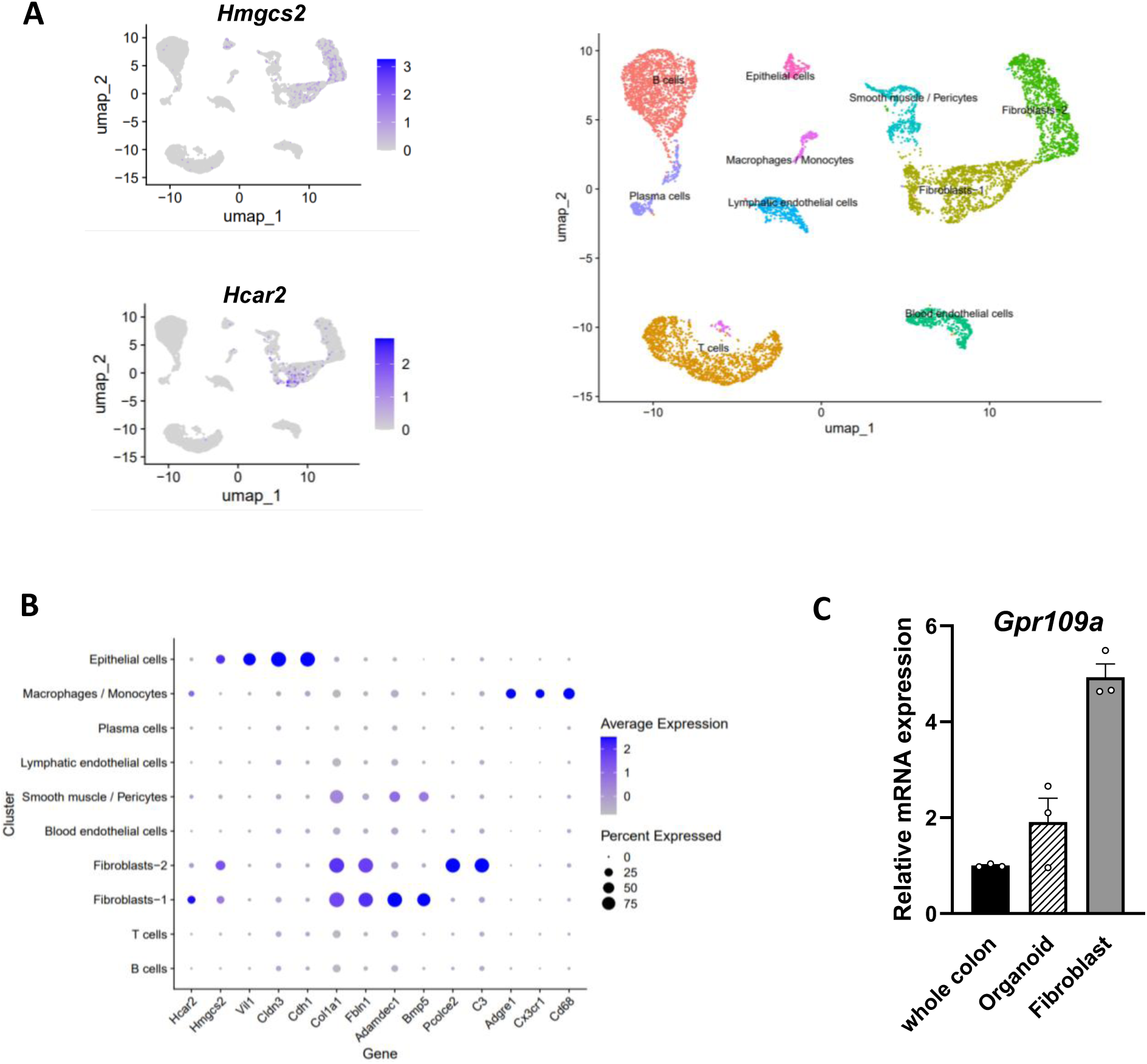
Cellular sources of GPR109A in the colon and its relationship to colonic ketogenesis. **(A, B)** Reanalysis of a public single-cell RNA-seq dataset. **(A)** UMAP plots showing cell populations colored by major cell type. **(B)** A dot plot displaying the expression profiles of *Hmgcs2*, *Gpr109a,* and selected marker genes. **(C)** *Gpr109a* expression in isolated colonic fibroblast cultures and epithelial organoids (n = 3).

## 3 DISCUSSION

In this study, we demonstrated that GPR109A exerts an indirect hepatoprotective effect through its expression in the colon by regulating intestinal TJ. KDs have been used as a therapeutic tool for over a century, although their effects on lipid metabolism remain debated. KDs can sometimes lead to elevated levels of LDL-C and total cholesterol and exacerbate fatty liver because of their high lipid content (7–10). In particular, studies in rodents have reported that high-fat KD contributes to the development of fatty liver and insulin resistance (28). Nevertheless, many cohort studies have reported the benefits of ketone bodies in improving neurological and metabolic disorders. Consequently, KDs are currently used in clinical settings when their benefits outweigh the associated risks. Accordingly, a comprehensive molecular understanding of both the beneficial and deleterious consequences of KDs is critical for their safe and effective application.

Our study confirmed that the high-fat KD induces hepatic inflammation and steatosis in mice. Furthermore, *Gpr109a^-/-^* mice exhibited more severe ectopic fat deposition in the liver, likely due to the upregulation of *de novo* lipogenesis. In addition, hepatic inflammation and fibrosis were further exacerbated in *Gpr109a^⁻/⁻^* mice, accompanied by an increased expression of the macrophage marker *Adgre1* and monocyte chemoattractant chemokine *Mcp1* in the liver. These findings indicate that GPR109A plays a critical role in maintaining hepatic homeostasis under KD conditions, in part by restraining macrophage recruitment and inflammation.

In contrast, during fasting conditions, while hepatic triglyceride levels were increased to the same levels comparable to those observed under KD-feeding conditions, *Tnf* expression in the liver remained unchanged in both WT and *Gpr109a^-/-^* mice. Moreover, no significant differences in hepatic triglycerides or inflammation were found between WT and *Gpr109a^-/-^*mice under fasting conditions. This discrepancy likely arises from the fundamentally distinct nature of these two ketogenic conditions: KD represents excessive lipid intake, whereas fasting reflects energy depletion, leading to divergent physiological adaptations. One such difference involves the intestinal barrier. Chronic exposure to a high-fat diet has been shown to impair barrier integrity (24). Previous studies have shown that activation of GPR109A, expressed in both epithelial and immune cells, by butyrate or nicotinic acid contributes to intestinal barrier maintenance in the DSS colitis model (17). Building upon this, we hypothesized that βHB–GPR109A signaling plays a crucial role in intestinal barrier integrity under high-fat KD conditions. Indeed, under KD but not fasting conditions, *Gpr109a^-/-^* mice exhibited reduced TJ markers specifically in the colon. Previous studies have also reported that GPR109A knockdown in Caco-2 cells suppressed sodium butyrate-induced Claudin-3 expression (29). Therefore, GPR109A may not act under conditions where intestinal barrier integrity remains intact, such as short-term fasting, but rather functions to counteract the deleterious effects of chronic high-fat intake on the gut epithelium. Intestinal barrier dysfunction allows translocation of PAMPs, which can drive hepatic inflammation and promote the progression of MASH (metabolic dysfunction-associated steatohepatitis). In our study, the expression of TLR4 signaling–related genes were upregulated in the liver of *Gpr109a^⁻/⁻^* mice, and treatment with antibiotics attenuated the hepatic inflammation observed in *Gpr109a^⁻/⁻^* mice under KD feeding. These results suggest that βHB–GPR109A signaling contributes to hepatic homeostasis under KD through indirect protection via the gut–liver axis.

Importantly, our findings suggest that GPR109A functions not merely as a receptor for βHB but as a sensor of the surrounding nutritional environment. GPR109A is also activated by butyrate, which is normally abundant in the gut. Thus, under normal chow conditions, luminal butyrate activates GPR109A. However, because the intestinal barrier remains intact, GPR109A-dependent protection is not manifested. In contrast, under KD conditions, the composition of the gut microbiota is altered, leading to a marked reduction in short-chain fatty acid production, including butyrate (approximately 2 mM) (12). Considering that the EC₅₀ of butyrate for GPR109A activation is approximately 1.6 mM (13), butyrate may barely activate the receptor, but its contribution is markedly reduced. In this context, βHB becomes the predominant ligand for GPR109A, effectively “switching” receptor activation, and mitigates the intestinal damage induced by the extremely high-fat composition of the KD. Short-term fasting also elevates intestinal ketone body levels; however, as fasting does not compromise barrier integrity, GPR109A-dependent protection is again not observed. Intermittent fasting, on the contrary, has been reported to negatively affect intestinal barrier function through repeated cycles of prolonged fasting and feeding despite its beneficial effects on weight control (30). Therefore, GPR109A may protect intestinal barrier integrity under intermittent fasting conditions. Long-term high-fat diet feeding is also known to disrupt the intestinal barrier (24). However, under commonly used approximately 60% fat diets, short-chain fatty acid levels drop dramatically (<2 mM) (31), yet glucose availability remains sufficient, and ketogenesis is minimal. Thus, under such high-fat diet conditions, the intestinal barrier is thought to deteriorate regardless of genotype, and GPR109A-mediated rescue may be unlikely. In addition to increased dietary fat intake in modern societies, supplements and drugs designed to inhibit lipid absorption in the small intestine have recently gained attention for their strong anti-obesity effects. While effective, these agents have been reported to adversely affect intestinal homeostasis (32, 33). Delivering ketone bodies or GPR109A agonists to the intestine may counteract such adverse effects, highlighting their potential therapeutic utility.

In recent years, it has been revealed that *Hmgcs2*, the rate-limiting enzyme of ketogenesis, is expressed not only in the liver but also in extrahepatic tissues such as the brain, kidney, and colon (25, 34). Previous studies have also demonstrated that colonic ketogenesis is essential for maintaining TJ organization (26). Therefore, we measured βHB levels in the colons of mice fed the KD and found that under diet-induced ketogenic conditions, tissue βHB levels were drastically elevated in addition to plasma levels. Recent studies have also reported that colonic ketogenesis increases circulating ketone body levels and that mice lacking intestinal ketone body-producing capacity exhibit increased intestinal permeability during colitis, raising the possibility that GPR109A may partially contribute to the this intestinal barrier integrity (26). Finally, to identify the colonic cell types that express *Gpr109a*, we reanalyzed a public single-cell RNA-seq dataset and confirmed its expression in epithelial cells and macrophages as previously reported. Notably, our analysis also indicated that *Gpr109a* is expressed in fibroblasts, suggesting a broader cellular distribution than previously thought. Recent studies have shown that fibroblasts actively contribute to epithelial homeostasis, including epithelial repair, through the secretion of various bioactive factors (35). Further studies using conditional knockout models or cell-type-specific ablation strategies will be essential to clarify the mechanisms underlying extrahepatic ketone body production and their functional interactions with GPCRs in the colon.

The role of GPR109A in lipid metabolism has also been widely studied. Since the discovery that nicotinic acid and subsequently βHB act as ligands for GPR109A (13, 18), the receptor was proposed to regulate lipolysis and LDL cholesterol levels (13). However, some studies report that GPR109A activation does not lead to a reduction in adipose tissue weight or plasma NEFA and cholesterol levels (36, 37), leaving some uncertainty regarding the role of GPR109A in lipid metabolism. In our study, acute βHB administration appeared to decrease circulating free fatty acids through GPR109A signaling, suggesting that GPR109A may modulate immediate lipolysis or fatty acid utilization. Nevertheless, *Gpr109a^-/-^* mice exhibited no significant differences in white adipose tissue mass or plasma NEFA concentrations under both KD feeding and fasting conditions. Consistently, clinical trials in patients with diabetes using GSK256073, a GPR109A agonist, also demonstrated that its effect on lipolysis was unfortunately diminished with prolonged treatment, likely due to the development of tolerance (38, 39). Together, these findings support the hypothesis that, under steady-state activation of GPR109A, its long-term effects on lipolysis may be masked by metabolic adaptation.

While our findings provide mechanistic insight into how GPR109A protects against KD-induced hepatic inflammation and steatosis, a few limitations should be acknowledged. While we demonstrated a suppression of hepatic inflammation by regulating intestinal TJ, we did not directly identify which GPR109A-expressing cell type mediates this effect. Future experiments using conditional knockout mice will be essential to address this question. Furthermore, we raise the possibility that ketone bodies synthesized in the intestinal epithelium may act in a paracrine manner to maintain intestinal barrier integrity. However, we cannot rule out the alternative possibility that βHB produced in the liver circulates through the bloodstream and contributes to these intestinal effects. Future experiments using organ-specific *Hmgcs2-*deficient mice, in combination with ketone body receptor*-*deficient mice, are required to clarify this issue. Another limitation relates to the translational relevance of our murine model. Mice and humans differ substantially in metabolic rate, lipid handling, and ketone body utilization, which can alter the physiological response to KDs. However, recent case reports in humans have described a MASH-like pathology during weight loss on a KD (10), underscoring that studies on gut–liver interactions under ketogenic conditions remain highly relevant.

In conclusion, we demonstrated that βHB-GPR109A signaling plays a crucial role in regulating the gut–liver axis, revealing a novel mechanism by which the body maintains metabolic homeostasis in response to ketone bodies. In addition, our study provides insights into the potential adverse effects of KDs and suggests that targeting GPR109A could provide a therapeutic strategy to protect intestinal barrier integrity and liver function under high-fat diet conditions.

## 4 MATERIALS AND METHODS

### 4.1 Animals and diet

Male C57BL/6J mice were purchased from Japan SLC (RRID: IMSR_JAX:000664, Shizuoka, Japan). *Gpr109a^−/−^* mice (C57BL/6J background) were generated using the CRISPR/Cas9 system (**Supplementary Figure 1**). Wild-type (WT) and *Gpr109a^−/−^* mice were originally derived from heterozygous intercrosses, and individual experiments included mice from at least three independent litters. Mice were housed in a conventional animal housing facility at 24°C under a 12 h/12 h light/dark cycle with *ad libitum* access to food and water and acclimated to the CLEA Rodent Diet (CE-2; CLEA Japan, Inc., Tokyo, Japan). For the ketogenic diet (KD) trial, 5-week-old mice were fed a normal diet (ND, CE-2) or a low-carbohydrate, high-fat diet with 90% kcal fat (KD, F3666, Bio-Serv., Flemington, NJ, USA) for 5weeks following a 1-week adaptation period, as previously described (12). The diet composition is presented in **Supplementary Table 1**. Antibiotic administration was conducted to deplete the gut microbiota, following a previously described method (40, 41). During the 5-week KD intervention, mice were provided with normal drinking water or water containing ampicillin sodium (0.5 mg/mL, FUJIFILM Wako Pure Chemical Corporation, Osaka, Japan), neomycin sulfate (0.5 mg/mL, FUJIFILM Wako Pure Chemical Corporation), metronidazole (0.5 mg/mL, FUJIFILM Wako Pure Chemical Corporation), and vancomycin hydrochloride (0.1 mg/mL, FUJIFILM Wako Pure Chemical Corporation,). For the fasting trial, 7-week-old mice were fasted for 24 h, as described previously (12). Body weight was measured once a week during long-term interventions. All mice were sacrificed using deep isoflurane-induced anesthesia, after which blood and tissue samples were collected.

### 4.2 Biochemical analyses

Plasma and hepatic levels of non-esterified fatty acids (NEFA), triglycerides, and total cholesterol were measured in mice using assay kits (LabAssay™ NEFA, triglyceride, and cholesterol; FUJIFILM Wako Pure Chemical Corporation) according to the manufacturer’s instructions. Blood glucose levels were measured using a portable glucometer (OneTouch^®^ Ultra^®^; LifeScan, Milpitas, CA, USA).

### 4.3 β-hydroxybutyrate (βHB) measurement

Plasma and intestinal βHB measurements were conducted following a previously described method with some modifications (12). Briefly, plasma samples and tissues were combined with cold acetonitrile containing an internal standard (sodium butyrate-^13^C_4_, Sigma-Aldrich, St. Louis, MO, USA), and the tissue samples were then homogenized. The samples were vortex-mixed for 30 s and then centrifuged at 10,000 × *g* for 5 min. The supernatant was subjected to liquid chromatography with tandem mass spectrometry (LC-MS/MS) analysis using an ultra-performance LC system (UPLC, Waters, Milford, MA, USA) equipped with an Acquity UPLC system coupled to a Waters Xevo TQD mass spectrometer (Waters). Samples were separated using an acetonitrile gradient in water containing 0.1% formic acid on an ACQUITY UPLC BEH C18 column (2.1 × 150 mm, 1.7 µm; Waters).

### 4.4 RNA isolation and quantitative reverse transcription (qRT)-PCR

Total RNA was extracted from animal tissues using the RNAiso Plus reagent (TAKARA, Shiga, Japan) in accordance with the manufacturer’s instructions. Complementary DNA (cDNA) was synthesized from the RNA templates using Moloney murine leukemia virus reverse transcriptase (Invitrogen, Thermo Fisher Scientific, Waltham, Massachusetts, USA). qRT-PCR was performed on a StepOne Real-Time PCR System (Applied Biosystems, Thermo Fisher Scientific, Waltham, Massachusetts, USA) with SYBR Premix Ex Taq II (TAKARA). The PCR conditions were as follows: 95°C for 30 s followed by 40 cycles of 95°C for 5 s, 58°C for 30 s, and 72°C for 1 min. Relative mRNA expression levels were quantified using the 2^-ΔΔCt^ method, with normalization to 18S rRNA as the housekeeping gene. Primer sequences are presented in **Supplementary Table 2**.

### 4.5 Histological analysis

Paraffin-embedded liver samples were sliced into 4-µm–thick sections. Immunofluorescence analysis was performed according to a standard protocol. Briefly, the paraffin-embedded cross-section was deparaffinized, fixed with 4% paraformaldehyde, permeabilized with 0.2% Triton X-100 (Sigma-Aldrich), and blocked with 1% bovine serum albumin (BSA) in PBS. Next, the sections were incubated with primary antibodies, followed by incubation with secondary antibodies conjugated with a fluorescent marker. Primary antibodies against F4/80 (1:1,000; Abcam, Cambridge, UK, ab6640), αSMA (1:300; Cell Signaling Technology, Danvers, MA, USA, 19245), and Claudin3 (1:50; Invitrogen, 34-1700) and secondary antibodies against goat anti-rat IgG antibody, biotinylated (1:150, BA-9400, Vector Laboratories, Newark, CA, USA) and goat anti-rabbit IgG antibody, biotinylated (1:150, BA-1000, Vector Laboratories) were used, respectively, and the nuclei were stained with DAPI (1:5000; Roche, Basel, Switzerland, 10236276001), as previously described (21). For hematoxylin-eosin (HE) staining, the paraffin-embedded sections fixed with 4% paraformaldehyde were stained with hematoxylin solution, followed by immersion in 70% ethanol for fixation, and then stained with the eosin solution. After being rinsed under running water, the sections underwent dehydration with 70% and 100% ethanol, and transparency with xylene. All sections were observed under a fluorescence and bright-field microscope (BZ-X710, Keyence Co., Osaka, Japan).

### 4.6 Flow cytometry

Hepatic cell isolation was performed according to a previously described protocol (21). Briefly, the liver was minced and treated with HBSS containing 3 mM CaCl₂, collagenase type I (1 mg/mL, FUJIFILM Wako Pure Chemical Corporation), and 1.5% BSA at 37°C for 2 h. The tissue was filtered through a 70-µm cell strainer to obtain single-cell suspensions. The suspensions were enriched using a Percoll density gradient (42) and then blocked with anti-CD16/CD32 (clone 93, BioLegend, San Diego, CA, USA) for 10 min at 4°C. Cells were subsequently stained with anti-CD45 (BV510, clone 30-F11, BioLegend), anti-Ly6C (BV711, clone HK1.4, BioLegend), anti-F4/80 (Alexa Fluor 488, clone BM8, BioLegend), anti-CX3CR1 (PE, clone SA011F11, BioLegend), and anti-CD11b (APC, clone M1/70, BioLegend) for 30 min at 4°C. The cells were next washed in FACS buffer consisting of PBS with 2% fetal bovine serum (FBS) and 2 mM EDTA. Cell sorting was carried out on FACSAria III or FACSMelody instruments (BD Biosciences, Franklin Lakes, New Jersey, USA), yielding populations with >95% purity.

### 4.7 Cell culture and intestinal organoid culture

All cell lines were cultured at 37°C under 5% CO_2_. **RAW264.7 cells** (mouse macrophage cell line; ATCC, Manassas, VA, USA) were cultured in Dulbecco’s Modified Eagle Medium (DMEM, FUJIFILM Wako Pure Chemical Corporation) supplemented with 10% FBS, (Gibco, Thermo Fisher Scientific, Waltham, MA, USA) and 1% penicillin–streptomycin (Gibco) (43). Cells were stimulated with βHB (1 and 10 mM, Sigma-Aldrich), MK-6892 (MedChemExpress LLC, Monmouth Junction, NJ, USA), or capric acid (C10:0; Nu-Chek Prep, Elysian, MN, USA) as a positive control, in the presence of lipopolysaccharide (LPS, 1 µg/mL; Sigma-Aldrich) for 10 h. After stimulation, cells were harvested for RNA isolation. Each *in vitro* experiment was conducted independently three times. **To generate colon organoids**, colon tissues were incubated in PBS solution containing 5 mM EDTA and rotated for 40 min at 4°C. Colonic crypts were isolated from the colonic segments by repetitive pipetting. The isolated crypts were embedded in Matrigel and maintained in human IntestiCult organoid growth medium (Veritas, Santa Clara, CA, USA) containing penicillin and streptomycin. The organoids were maintained at 37°C in 5% CO_2_, and the medium was changed every other day. After 3 days of culture, the organoids were harvested for RNA isolation. **Primary culture of colonic fibroblasts** was performed with minor modifications to previously described methods (44). Colon tissue was incubated in PBS containing 20 mM EDTA at 37°C for 20 min. After a 1-min vortex, the EDTA/PBS solution was replaced with ice-cold PBS, and this washing step was repeated twice. The muscularis externa was then removed under a stereomicroscope, and the tissue was further vortexed in ice-cold PBS until the mucosa became translucent. The tissue was subsequently digested in an enzyme solution containing collagenase type II (1.5 mg/mL, Worthington, Lakewood, NJ, USA) and dispase (1 mg/mL, Gibco) for 20 min at 37°C with intermittent vigorous vortexing. The digest was passed through a 70-µm cell strainer to obtain dispersed cells, and enzymatic activity was quenched by adding DMEM supplemented with 40% FBS. Cells were pelleted by centrifugation at 350 × *g* for 5 min and seeded at 4 × 10⁵ cells per well in 48-well plates. Total RNA was collected after 48 h of culture.

### 4.8 Single-cell RNA sequencing (scRNA-Seq) data reanalysis

Publicly available scRNA-Seq data were obtained from the Gene Expression Omnibus under accession number GSE264408, originally published previously (27). For the present analysis, we used the dataset derived from a healthy mouse colon sample (Healthy3). **Quality control and filtering:** The raw barcodes.tsv, features.tsv, and matrix.mtx files were imported into R (version 4.5.1) and processed using the Seurat package. Cells with <1,000 or >5,000 detected genes, total counts >30,000, or mitochondrial gene expression >10% were excluded. **Normalization and dimensionality reduction:** Normalized expression values were calculated using SCTransform. Principal component analysis (PCA) was then performed on the scaled data, and the number of principal components (PCs) used for downstream analyses was determined using an elbow plot. The first 10 PCs were selected for downstream analyses. **Clustering:** A K-nearest neighbor graph was constructed using the selected PCs, and cells were clustered using the Louvain community detection algorithm (resolution = 0.1). Clusters were visualized using uniform manifold approximation and projection (UMAP) embeddings. Cluster identities were assigned based on the expression of canonical marker genes (**Supplementary Figure 5**).

### 4.9 Statistical analysis

All data are presented as the mean ± standard error of the mean (SEM). Statistical analyses were conducted using GraphPad Prism (GraphPad Software Inc., La Jolla, CA, USA, RRID: SCR_002798). Data normality was assessed using the Shapiro–Wilk test. For comparisons between two groups, either a two-tailed Student’s t-test or Mann–Whitney U test was used as appropriate. For experiments involving two independent variables, statistical significance was evaluated using two-way analysis of variance (ANOVA) followed by Sidak’s multiple comparisons test. For cell-based assays involving comparisons with a single control group, Dunnett’s multiple comparisons test was applied. A P-value <0.05 was considered statistically significant.

### 4.10 Study approval

All experimental procedures involving mice were performed in accordance with the guidelines of the Committee on the Ethics of Animal Experiments of the Kyoto University Animal Experimentation Committee. Permission for the use of animals in this study was granted by Kyoto University under protocol number Lif-K26002. All efforts were made to minimize animal suffering.

### 4.11 Data availability statement

All data generated or analyzed during this study are included in this published article and its Supplementary files or are available from the corresponding authors upon reasonable request.

## AUTHORSHIP CONTRIBUTIONS

**A.N.** was responsible for conceptualization and performed experiments, interpreted data, and wrote the paper. **S.N.** performed experiments, interpreted data, and reviewed the paper. **R.B.** performed the experiments and wrote the paper. **M.Y.** performed experiments and reviewed the paper. **R.O.-K.** performed the experiments and interpreted data. **T.I.** interpreted data and reviewed the paper. **N.S.** performed the experiments and interpreted the data. **I.K.** supervised the project, interpreted data, and wrote the paper. **A.N.** and **I.K.** are the guarantors of this work and, as such, have full access to all the data in the study and take responsibility for the integrity of the data and the accuracy of the data analysis. All authors read and approved the final manuscript.

## REFERENCES

1. Puchalska P, and Crawford PA. Multi-dimensional Roles of Ketone Bodies in Fuel Metabolism, Signaling, and Therapeutics. Cell Metab. 2017;25(2):262–84.

2. Kashiwaya Y, et al. D-beta-hydroxybutyrate protects neurons in models of Alzheimer’s and Parkinson’s disease. Proc Natl Acad Sci. 2000;97(10):5440–4.

3. Dmitrieva-Posocco O, et al. β-Hydroxybutyrate suppresses colorectal cancer. Nature. 2022;605(7908):160–5.

4. Huttenlocher PR, et al. Medium-chain triglycerides as a therapy for intractable childhood epilepsy. Neurology. 1971;21(11):1097–103.

5. Kossoff EH, and Dorward JL. The modified Atkins diet. Epilepsia. 2008;49 Suppl 8:37–41.

6. Pfeifer HH, and Thiele EA. Low-glycemic-index treatment: a liberalized ketogenic diet for treatment of intractable epilepsy. Neurology. 2005;65(11):1810–2.

7. Batch JT, et al. Advantages and Disadvantages of the Ketogenic Diet: A Review Article. Cureus. 2020;12(8):e9639.

8. Bueno NB, et al. Very-low-carbohydrate ketogenic diet v. low-fat diet for long-term weight loss: a meta-analysis of randomised controlled trials. Br J Nutr. 2013;110(7):1178–87.

9. Nordmann AJ, et al. Effects of low-carbohydrate vs low-fat diets on weight loss and cardiovascular risk factors: a meta-analysis of randomized controlled trials. Arch Intern Med. 2006;166(3):285–93.

10. Anekwe CV, et al. enic Diet-induced Elevated Cholesterol, Elevated Liver Enzymes and Potential Non-alcoholic Fatty Liver Disease. Cureus. 2020;12(1):e6605.

11. Kimura I, et al. Short-chain fatty acids and ketones directly regulate sympathetic nervous system via G protein-coupled receptor 41 (GPR41). Proc Natl Acad Sci U S A. 2011;108(19):8030–5.

12. Miyamoto J, et al. Ketone body receptor GPR43 regulates lipid metabolism under ketogenic conditions. Proc Natl Acad Sci U S A. 2019;116(47):23813–21.

13. Taggart AK, et al. (D)-beta-Hydroxybutyrate inhibits adipocyte lipolysis via the nicotinic acid receptor PUMA-G. J Biol Chem. 2005;280(29):26649–52.

14. Kimura I, et al. Free Fatty Acid Receptors in Health and Disease. Physiol Rev. 2020;100(1):171–210.

15. Benyó Z, et al. GPR109A (PUMA-G/HM74A) mediates nicotinic acid-induced flushing. J Clin Invest. 2005;115(12):3634–40.

16. Shenol A, et al. Multiple recent HCAR2 structures demonstrate a highly dynamic ligand binding and G protein activation mode. Nat Commun. 2024;15(1):5364.

17. Singh N, et al. Activation of Gpr109a, receptor for niacin and the commensal metabolite butyrate, suppresses colonic inflammation and carcinogenesis. Immunity. 2014;40(1):128–39.

18. Tunaru S, et al. PUMA-G and HM74 are receptors for nicotinic acid and mediate its anti-lipolytic effect. Nat Med. 2003;9(3):352–5.

19. Offermanns S. The nicotinic acid receptor GPR109A (HM74A or PUMA-G) as a new therapeutic target. Trends Pharmacol Sci. 2006;27(7):384–90.

20. Thangaraju M, et al. GPR109A is a G-protein-coupled receptor for the bacterial fermentation product butyrate and functions as a tumor suppressor in colon. Cancer Res. 2009;69(7):2826–32.

21. Ohue-Kitano R, et al. Medium-chain fatty acids suppress lipotoxicity-induced hepatic fibrosis via the immunomodulating receptor GPR84. JCI Insight. 2023;8(2).

22. Zheng L, et al. Microbial-Derived Butyrate Promotes Epithelial Barrier Function through IL-10 Receptor-Dependent Repression of Claudin-2. J Immunol. 2017;199(8):2976–84.

23. Chen G, et al. Sodium Butyrate Inhibits Inflammation and Maintains Epithelium Barrier Integrity in a TNBS-induced Inflammatory Bowel Disease Mice Model. EBioMedicine. 2018;30:317–25.

24. Kawano Y, et al. Colonic Pro-inflammatory Macrophages Cause Insulin Resistance in an Intestinal Ccl2/Ccr2-Dependent Manner. Cell Metab. 2016;24(2):295–310.

25. Camarero N, et al. Ketogenic HMGCS2 Is a c-Myc target gene expressed in differentiated cells of human colonic epithelium and down-regulated in colon cancer. Mol Cancer Res. 2006;4(9):645–53.

26. Bass K, et al. Colonic ketogenesis, a microbiota-regulated process, contributes to blood ketones and protects against colitis in mice. Biochem J. 2024;481(4):295–312.

27. Hong D, et al. Integrative analysis of single-cell RNA-seq and gut microbiome metabarcoding data elucidates macrophage dysfunction in mice with DSS-induced ulcerative colitis. Commun Biol. 2024;7(1):731.

28. Jornayvaz FR, et al. A high-fat, ketogenic diet causes hepatic insulin resistance in mice, despite increasing energy expenditure and preventing weight gain. Am J Physiol Endocrinol Metab. 2010;299(5):E808–15.

29. Feng W, et al. Sodium Butyrate Attenuates Diarrhea in Weaned Piglets and Promotes Tight Junction Protein Expression in Colon in a GPR109A-Dependent Manner. Cell Physiol Biochem. 2018;47(4):1617–29.

30. Liang Y, et al. Maternal intermittent fasting in mice disrupts the intestinal barrier leading to metabolic disorder in adult offspring. Commun Biol. 2023;6(1):30.

31. Shimizu H, et al. Dietary short-chain fatty acid intake improves the hepatic metabolic condition via FFAR3. Sci Rep. 2019;9(1):16574.

32. Morales P, et al. Impact of Dietary Lipids on Colonic Function and Microbiota: An Experimental Approach Involving Orlistat-Induced Fat Malabsorption in Human Volunteers. Clin Transl Gastroenterol. 2016;7(4):e161.

33. Katimbwa DA, et al. Orlistat, a competitive lipase inhibitor used as an antiobesity remedy, enhances inflammatory reactions in the intestine. Applied Biological Chemistry. 2022;65(1):47.

34. Tomita I, et al. SGLT2 Inhibition Mediates Protection from Diabetic Kidney Disease by Promoting Ketone Body-Induced mTORC1 Inhibition. Cell Metab. 2020;32(3):404–19.e6.

35. Chalkidi N, et al. Fibroblasts in intestinal homeostasis, damage, and repair. Front Immunol. 2022;13:924866.

36. Geisler CE, et al. The Role of GPR109a Signaling in Niacin Induced Effects on Fed and Fasted Hepatic Metabolism. Int J Mol Sci. 2021;22(8).

37. Lauring B, et al. Niacin lipid efficacy is independent of both the niacin receptor GPR109A and free fatty acid suppression. Sci Transl Med. 2012;4(148):148ra15.

38. Dobbins RL, et al. GSK256073, a selective agonist of G-protein coupled receptor 109A (GPR109A) reduces serum glucose in subjects with type 2 diabetes mellitus. Diabetes Obes Metab. 2013;15(11):1013–21.

39. Dobbins R, et al. GSK256073 acutely regulates NEFA levels via HCA2 agonism but does not achieve durable glycaemic control in type 2 diabetes. A randomised trial. Eur J Pharmacol. 2015;755:95–101.

40. Igarashi M, et al. Intestinal GPR119 activation by microbiota-derived metabolites impacts feeding behavior and energy metabolism. Mol Metab. 2023;67:101649.

41. Haneishi Y, et al. Polyunsaturated fatty acids-rich dietary lipid prevents high fat diet-induced obesity in mice. Sci Rep. 2023;13(1):5556.

42. Lynch RW, et al. An efficient method to isolate Kupffer cells eliminating endothelial cell contamination and selective bias. J Leukoc Biol. 2018;104(3):579–86.

43. Nakajima A, et al. The short chain fatty acid receptor GPR43 regulates inflammatory signals in adipose tissue M2-type macrophages. PLoS One. 2017;12(7):e0179696.

44. Lee RF, et al. A Coculture System for Modeling Intestinal Epithelial-Fibroblast Crosstalk. Methods Mol Biol. 2025;2951:19–34.

