## Supplemental Materials for "Ketone-body receptor GPR109A suppresses hepatic inflammation via gut–liver axis regulation"

### Supplementary Figure 1

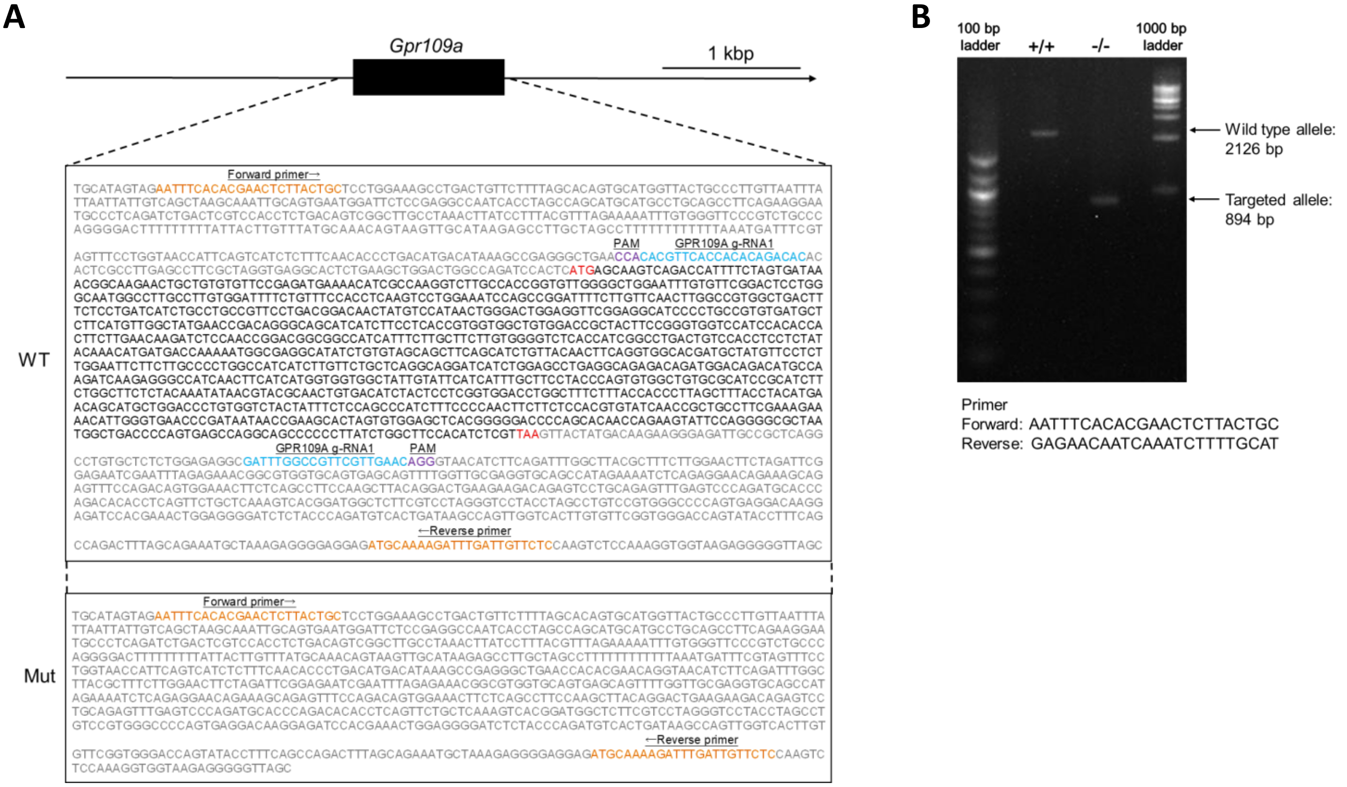


**Supplementary Figure 1.** Generation of *Gpr109a* deficient (*Gpr109a*$^{-/-}$) mice. **(A)** The coding exon regions are identified using black boxes and words. Guide RNA (gRNA) and protospacer adjacent motif (PAM) are indicated in light blue and purple, respectively. **(B)** Mouse genotypes were determined by PCR using the indicated primers to detect WT and mutant alleles of *Gpr109a*.

**Supplementary Figure 2**

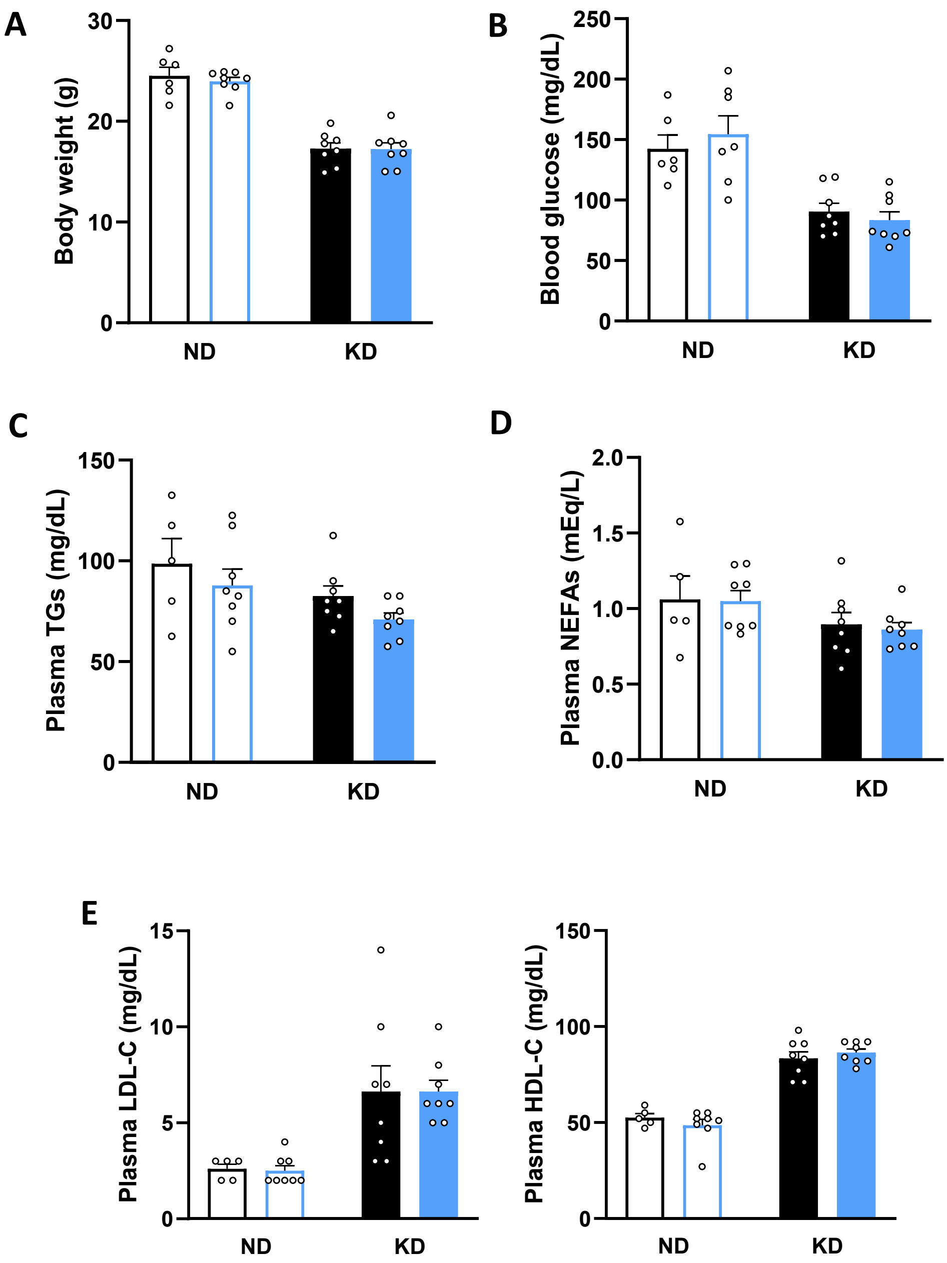


**Supplementary Figure 2.** Metabolic parameters in WT and *Gpr109a*$^{-/-}$ mice fed a diet-induced ketogenic diet. **(A)** Body weight at 11 weeks prior to sacrifice (n = 6–8). **(B–E)** Blood glucose (B, n = 6–8), plasma triglycerides (C, n = 5–8), plasma NEFAs, (D, n = 5–8), plasma LDL- and HDL-cholesterol (E, n = 5–8). two-way ANOVA with Sidak's multiple comparisons test: A–E. All data are presented as the mean ± SEM.

#### Supplementary Figure 3

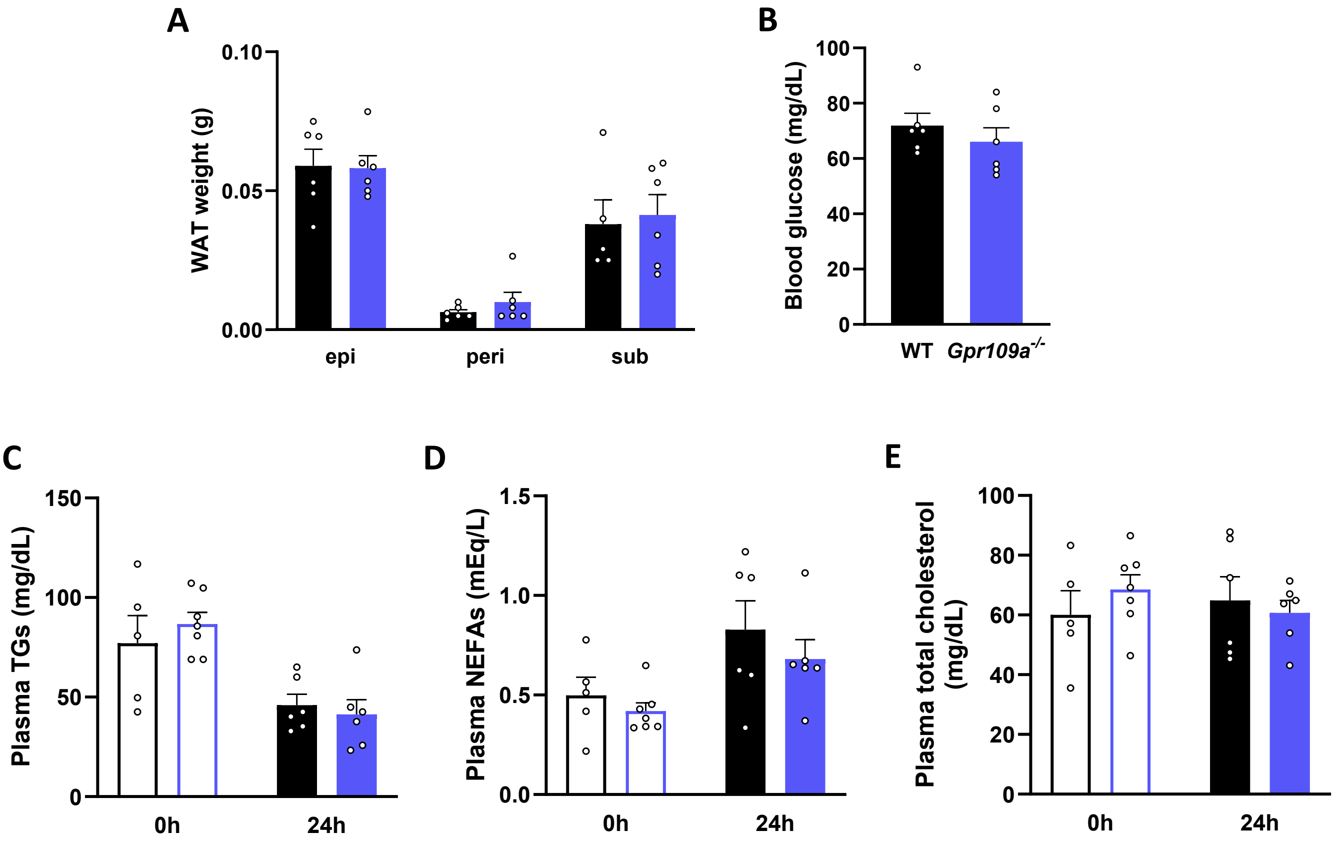


**Supplementary Figure 3.** Metabolic parameters in WT and *Gpr109a*$^{-/-}$ mice under fasting-induced ketogenic diet. **(A)** White adipose tissue weight (n = 5–6). **(B–E)** Blood glucose (B, n = 6) and plasma triglycerides (C, n = 5–7), plasma NEFAs, (D, n = 5–7) plasma total cholesterol (E, n = 5–7). Student's t-test: A, B; two-way ANOVA with Sidak's multiple comparisons test: C–E. All data are presented as the mean ± SEM.

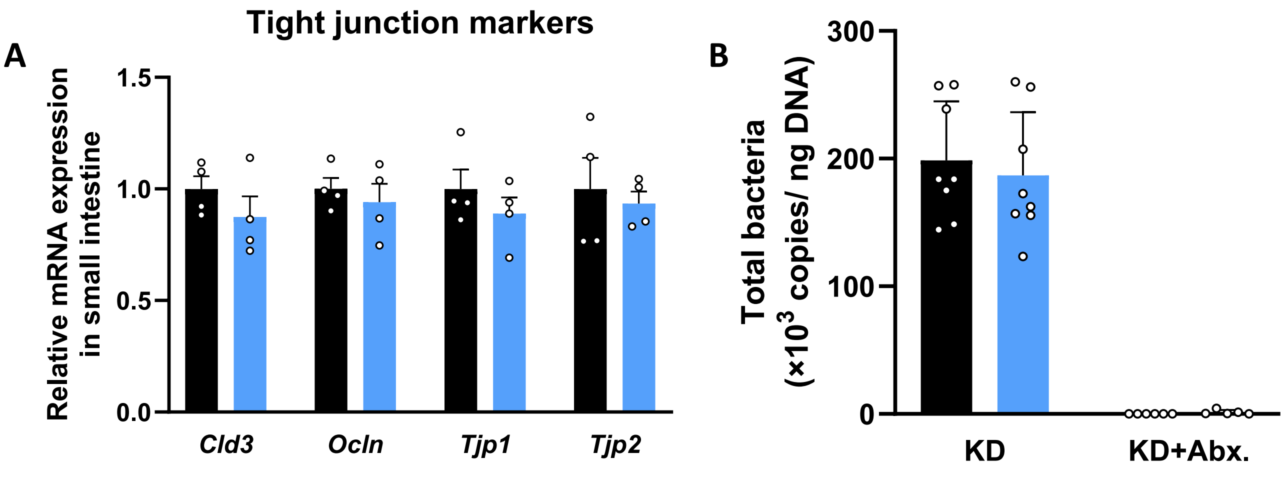


**Supplementary Figure 4.** GPR109A functions in the intestine. **(A)** mRNA expression levels of tight junction-related genes in the small intestine of WT and *Gpr109a*$^{-/-}$ mice under diet-induced ketogenic conditions (n = 4). **(B)** Copy numbers of gut microbiota determined by qRT-PCR (n = 5–8). Mann–Whitney U test: A. All data are presented as the mean ± SEM.

#### Supplementary Figure 5

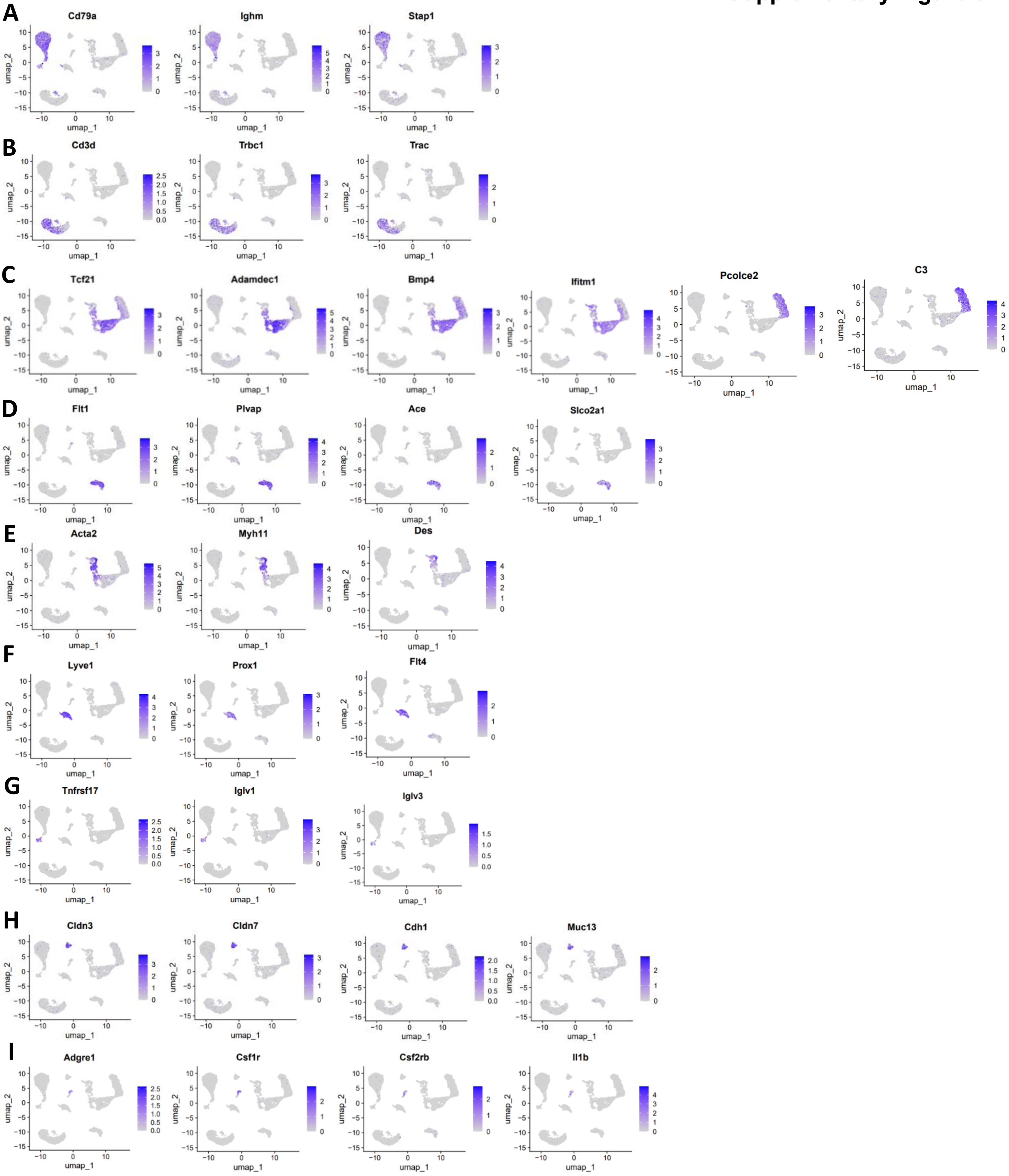


**Supplementary Figure 5.** Single-cell RNA-seq data were reanalyzed, and representative marker genes for each cell cluster are listed. **(A)** B cells. **(B)** T cells. **(C)** Fibroblasts. **(D)** Blood endothelial cells. **(E)** Smooth muscle cells/pericytes. **(F)** Lymphatic endothelial cells. **(G)** Plasma cells. **(H)** Macrophages/monocytes. **(I)** Epithelial cells.

**Supplementary Table 1.** Composition of the ketogenic diet (Product #F3666, Ketogenic Diet, AIN-76A Modified, High Fat, Paste).

| Proximate Profile | | |
|---|---|---|
| Protein | % | 8.6 |
| Fat | % | 75.1 |
| Fiber | % | 4.8 |
| Ash | % | 3.0 |
| Moisture | % | <10 |
| Carbohydrate | % | 3.2 |

| Caloric Profile | | |
|---|---|---|
| Protein | kcal/ gm | 0.34 |
| Fat | kcal/ gm | 6.76 |
| Carbohydrate | kcal/ gm | 0.13 |
| Total | | 7.24 |

| Amino Acids | | |
|---|---|---|
| Alanine | gm/kg | 2.3 |
| Arginine | gm/kg | 3.1 |
| Aspartic Acid | gm/kg | 5.5 |
| Cystine | gm/kg | 0.3 |
| Glutamic Acid | gm/kg | 17.3 |
| Glycine | gm/kg | 2.1 |
| Histidine | gm/kg | 2.3 |
| Isoleucine | gm/kg | 4.7 |
| Leucine | gm/kg | 7.1 |
| Lysine | gm/kg | 6.3 |
| Methionine | gm/kg | 2.2 |
| Phenylalanine | gm/kg | 3.8 |
| Proline | gm/kg | 8.7 |
| Serine | gm/kg | 4.8 |
| Threonine | gm/kg | 3.7 |
| Tryptophan | gm/kg | 1.0 |
| Tyrosine | gm/kg | 4.8 |
| Valine | gm/kg | 5.5 |

| Carbohydrates | | |
|---|---|---|
| Monosaccharides | gm/kg | 7.0 |
| Disaccharides | gm/kg | 24.9 |
| Polysaccharides | gm/kg | 0.0 |

| Fatty Acids | | |
|---|---|---|
| C18:2 Linoleic | gm/kg | 115 |
| C18:3 Linolenic | gm/kg | 6.7 |
| Total Saturated | gm/kg | 303 |
| Total Monounsaturated | gm/kg | 288 |
| Total Polyunsaturated | gm/kg | 122 |

| Minerals | | |
|---|---|---|
| Calcium | gm/kg | 5.7 |
| Chloride | gm/kg | 1.7 |
| Copper | mg/kg | 6.6 |
| Chromium | mg/kg | 2.2 |
| Fluoride | mg/kg | 0.0 |
| Iodine | mg/kg | 0.2 |
| Iron | mg/kg | 38.7 |
| Magnesium | gm/kg | 0.56 |
| Manganese | mg/kg | 63.6 |
| Phosphorus | gm/kg | 4.9 |
| Potassium | gm/kg | 3.9 |
| Selenium | mg/kg | 0.19 |
| Sodium | mg/kg | 1128 |
| Sulfur | mg/kg | 366 |
| Zinc | mg/kg | 36.0 |

| Vitamins | | |
|---|---|---|
| Ascorbic Acid | mg/kg | 0.0 |
| Biotin | mg/kg | 0.42 |
| Choline | mg/kg | 274 |
| Folic Acid | mg/kg | 4.2 |
| Niacin | mg/kg | 62.8 |
| Pantothenic Acid | mg/kg | 30.9 |
| Pyridoxine | mg/kg | 12.1 |
| Riboflavin | mg/kg | 12.6 |
| Thiamin | mg/kg | 11.2 |
| Vitamin A | IU/kg | 15500 |
| Vitamin $B_{12}$ | mcg/kg | 21 |
| Vitamin $D_3$ | IU/kg | 2090 |
| Vitamin E | IU/kg | 244 |
| Vitamin $K_3$(Menadione) | mg/kg | 2.2 |

**Ingredients**

Lard, Butter, Corn Oil, Casein, Cellulose
Mineral Mix, Vitamin Mix, Dextrose

**Supplementary Table 2.** Primer sequences used for quantitative real-time PCR (qPCR).

| gene | Forward | Reverse |
|---|---|---|
| *18s* | CTCAACACGGGAAACCTCAC | AGACAAATCGCTCCACCAAC |
| *Dgat2* | TCTTCTGGACCCATCGGCCCCAGGA | AGTGGCAATGCTATCATCATCGT |
| *Srebf1* | GGAGCCATGGATTGCACATT | GGCCCGGGAAGTCACTGT |
| *Fabp4* | GATGCCTTTGTGCGAACCTGG | CTGTCGTCTGCGGTGATTTC |
| *Acaca* | CTTCCTGACAAACGAGTCTGG | CTGCCGAAACATCTCTGGGA |
| *Pparg2* | GCTGTTATGGGTGAAACTCTGG | TTCTTGTGAAGTGCTCATAGGC |
| *Scd1* | GCAAGCTCTACACCTGCCTCTT | CGTGCCTTGTAAGTTCTGTGGC |
| *Lpl* | CTGCTGGCGTAGCAGGAAGT | GCTGGAAAGTGCCTCCATTG |
| *Fatp1* | CATCCGTCTGGTCAAGGTCA | ACGCTGTGGGCAATCTTCTT |
| *Fas* | TTGCCCGAGTCAGAGAACCT | GTCCATTGTGTGTGCCTGCT |
| *Cd36* | TGGCAAAGAACAGCAGCAAA | GACAGTGAAGGCTCAAAGATGG |
| *Gpr109a* | CCGTTCCTGACGGACAACTA | ATGCCTCGCCATTTTTGGTC |
| *Tnf* | GGCAGGTCTACTTTGGAGTC | TCGAGGCTCCAGTGAATTCG |
| *Col1a* | CCTCAGGGTATTGCTGGACAAC | ACCACTTGATCCAGAAGGACCTT |
| *Adgre1* | GATGTGGAGGATGGGAGATG | ACAGCAGGAAGGTGGCTATG |
| *Tlr4* | CCGCTCTGGCATCATCTTC | TGTTCTTCCTCTGCTGTTTGCTC |
| *Il1b* | GCCCATCCTCTGTGACTCAT | AGGCCACAGGTATTTTGTCG |
| *Cldn3* | CAGTGTACCAACTGCGTACAAGAC | ACCGGTACTAAGGTGAGCAGAG |
| *Ocln* | GGACCCTGACCACTATGAAACAGACTAC | ATAGGTGGATATTCCCTGACCCAGTC |
| *Tjp1* | TGGGAACAGCACACAGTGAC | GCTGGCCCTCCTTTTAACAC |
| *Tjp2* | GACCTCAATCGTCATCTCAGATGT | GCTGCGGAAACTTCTGCCATCAAA |
